# Double mutant of chymotrypsin inhibitor 2 stabilized through increased conformational entropy

**DOI:** 10.1101/2021.11.18.469114

**Authors:** Yulian Gavrilov, Felix Kümmerer, Simone Orioli, Andreas Prestel, Kresten Lindorff-Larsen, Kaare Teilum

## Abstract

The conformational heterogeneity of a folded protein can affect both its function but also stability and folding. We recently discovered and characterized a stabilized double mutant (L49I/I57V) of the protein CI2 and showed that state-of-the-art prediction methods could not predict the increased stability relative to the wild-type protein. Here we have examined whether changed native state dynamics, and resulting entropy changes, can explain the stability changes in the double mutant protein, as well as the two single mutant forms. We have combined NMR relaxation measurements of the ps-ns dynamics of amide groups in the backbone and the methyl groups in the side-chains with molecular dynamics simulations to quantify the native state dynamics. The NMR experiments reveal that the mutations have different effects on the conformational flexibility of CI2: A reduction in conformational dynamics (and entropy) of the native state of L49I variant correlates with its decreased stability, while increased dynamics of the I57V and L49I/I57V variants correlates with their increased stability. These findings suggest that explicitly accounting for changes in native state entropy might be needed to improve the predictions of the effect of mutations on protein stability.

## Introduction

Controlling protein stability is an important challenge in biotechnology and biopharmaceutical applications. The ability to predict how protein mutations decrease stability can be used in clinical genetics to interpret gene or genome sequencing data.^1^ Similarly, improving the stability of enzymes^2^ or protein pharmaceuticals^3^ by mutations may improve shelf life and tolerance to heat. The stability of a protein depends on interactions and dynamics of both the native and denatured states of the protein.^4^

The conformational dynamics of the native state of a protein contributes to its conformational stability. If the dynamics of the native state is increased without significantly affecting the intermolecular interactions it will lead to a more stable protein.^5,6^ Typically, however, the increased dynamics is counteracted by fewer stabilizing interactions through entropy-enthalpy compensation in the more dynamic molecule.^7,8^ In some cases evolution has circumvented this problem. Thus, in hyperthermophilic proteins Lys is preferred over Arg compared to the mesophilic homologues. This is attributed to high residual entropy of the Lys side-chain relative to Arg in folded protein structures.^9^ However, even small local changes in a protein structure induced by point mutations can affect protein conformational equilibria, dynamics and its interactions.^10,11^ NMR spectroscopy, in combination with computational methods such as molecular dynamics simulations, has successfully been applied to analyze the effect of mutations on conformational dynamics of various protein systems.^8,12–17^ Here we apply these complementary techniques to study the small 7.3 kDa protein, chymotrypsin inhibitor 2 (CI2) from barley seeds,^18^ which has been extensively used in both experimental and computational studies.^19–24^ The conformational dynamics of the polypeptide backbone and the side-chains of CI2 on the ps-ns time-scales have been studied by NMR spectroscopy.^25^ Previously, it was demonstrated that five single point-mutations in the hydrophobic core of CI2 (L49V, L49A, I57V, V9A, and V47A) all lead to a similar yet small increase in backbone and side-chain flexibility throughout the structure. These (and other) core mutations only have a minor effect on the inhibitory function of CI2.^26^

According to experimental^19^ and computational studies^27,28^ residues L49 and I57 are both important for formation of the CI2 folding nucleus and for the stability of the folded state as they participate in a central network of interactions.^25–31^ L49 (together with R48) is a key energetic linchpin connecting CI2’s hydrophobic core to its inhibtory loop.^27,30^ Furthermore, L49 and I57 are involved in a network of interactions that dissipate the structural strain occurring when CI2 binds to its substrate.^31^ This network includes residue I37 in the inhibitory loop. The neighboring residues, T36 and V38, point their side chains toward the highly conserved residues of the inhibitor loop, R48 and F50. Between these residues L49 forms part of the hydrophobic core together with A16 and I57.^31^

We recently showed that L49I destabilizes the folded state of CI2 (ΔΔ*G*_f_ = 3.4±0.1 kJ/mol), while I57V stabilizes it slightly (ΔΔ*G*_f_ = −2.2±0.1 kJ/mol).^32^ Notably, the double mutant (L49I/I57V) is significantly more stable than either of the single mutants (ΔΔ*G*_f_ = −3.8±0.1 kJ/mol; Figure 1). This synergistic stabilizing effect of the double mutant is unexpected and could not be predicted using commonly applied methods such as FoldX^33^ and Rosetta.^34^ Failure to properly capture long range structural changes and changes in the dynamic properties of the protein may contribute to the unsuccessful attempt to predicting the effect of the double mutation.

**Figure 1.**
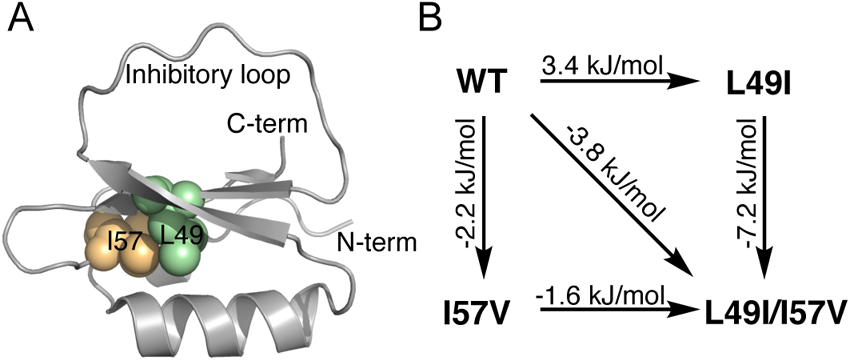
**(A)** Overview of the positions of the mutated sites in CI2 (spacefilling). (B) Mutation cycle labelled with experimental ΔΔ*G*_f_ values at 25 °C.

Here, we asked if the change in stability of the L49I/I57V double mutant can, at least in part, be explained by changes in the dynamics of CI2. We used NMR spectroscopy to experimentally assess the changes in fast dynamic of the three CI2 variants L49I, I57V and L49I/I57V relative to the wild type. To complement the experimental data, we also performed molecular dynamics (MD) simulations and reweighted the obtained computational dynamic ensembles using experimental data. We found that I57V and L49I/I57V increases the dynamics of CI2 whereas L49I decreases the dynamics. The corresponding change in conformational entropy scales linearly with the change in conformational stability. Our findings suggest that explicitly accounting for changes in native state entropy might be needed to improve the predictions of the effect of mutations on protein stability, which may have implications for the interpretation of protein engineering and sequencing data.

## Materials and methods

### Protein production and sample preparation

We expressed wild-type CI2 (UniProt: P01053, residues 22–84 with an additional N-terminal Met) and the three variants L49I, I57V, and L49I/I57V using previously described DNA constructs.^32^ *E. coli* BL21(DE3) carrying the expression plasmid was grown in M9 minimal medium at 37 ºC to OD_600_=0.6 before induction with 0.4 mM isopropyl b-D-1-thiogalactopyranoside (IPTG). Four different isotope compositions of the M9 media were used. i) ^15^NH_4_Cl as the sole nitrogen source, ii) ^13^C_6_-Glc as the sole carbon source and ^15^NH_4_Cl as the sole nitrogen source, iii) ^13^C_6_-Glc as the sole carbon source and ^15^NH_4_Cl as the sole nitrogen source in 50% D_2_O and iv) 10% ^13^C_6_-Glc / 90% unlabeled Glc as carbon source and ^15^NH_4_Cl as the sole nitrogen source. The growth was continued overnight at 20 ºC. Harvested cells were resuspended in 25mM Tris HCl, pH 8 with 1 mM of EDTA. Cell lysis was performed with two freeze-thaw cycles. Insoluble material was removed by centrifugation at 30000 g for 20 minutes at 5 ºC.

Nucleic acids were precipitated with 2% polyethylene imine and removed by centrifugation at 20000 g for 15 min at 5 ºC. CI2 was precipitated by adding ammonium sulfate to 70% saturation and centrifuging at 20000 g for 15 min at 5 ºC. The ammonium sulfate precipitate was resuspended in 10 mM ammonium bicarbonate and pure CI2 was obtained by SEC on a Superdex 75 26/600 in 10 mM ammonium bicarbonate.

### NMR Experiments

All NMR experiments were performed at constant temperature (25 ºC) and buffer conditions (50 mM MES, pH 6.26, 5 % D_2_O and 0.02 % NaN_3_). ^15^N-HSQC, HNCO, HN(CA)CO, HNCA, HNCACB, CBCA(CO)NH spectra were acquired and used for backbone chemical shift (CS) assignments. The methyl chemical shifts were assigned using ^13^C-HSQC, C(CO)NH and HC(CO)NH spectra. These spectra were acquired for all studied CI2 variants on a Varian DD2 800 MHz spectrometer equipped with a room temperature probe. The triple resonance spectra were recorded with non-uniform sampling at 25% and reconstructed by the compressed sensing method iterative soft thresholding using the software qMDD.^35^ We stereospecifically assigned the methyl groups of the valine and leucine side chain from constant time ^13^C-HSQC recorded on the sample expressed in the presence of 10% ^13^C_6_-Glc.^36^

Backbone dynamics was probed with a series of ^15^N *R*_1_, *R*_2_ and heteronuclear ^1^H-^15^N NOE relaxation experiments recorded on Bruker Avance III HD 600 and 750 MHz spectrometers cryoprobes using ~1.5 mM ^15^N-labeled protein samples. Relaxation delays of 10, 200, 400, 600, 800, 1000 ms were chosen for *R*_1_ and 16.96, 84.8, 169.6, 254.4, 339.2, 424.8 ms for *R*_2_, respectively. The ^1^H-^15^N NOE experiments were recorded with a 5 s interscan delay. All experiments were repeated twice, processed with nmrPipe^37^ and analyzed in CcpNmr.^38^ We performed model-free analysis^39,40^ for the calculation of backbone order parameters *S*^2^_*NH*_ using the software *relax*.^41^ Values of *R*_*1*_, *R*_*2*_ and ^1^H-^15^N NOE for each amide group can be fitted to model-free models with up to six parameters.^42^ Our initial analysis showed that fitting to models with more than three parameters resulted in unrealistic values of the order parameters. In the final analysis of the backbone NMR data, we therefore excluded models with more than three parameters.

Side chain ^2^H relaxation experiments were performed on ~0.7 mM ^15^N/^13^C/^2^H_50%_-labeled protein samples using pulse sequences described elsewhere.^43–45^ The relaxation data were acquired on Bruker Avance III HD 600 and 750 MHz spectrometers using following delays for *R*_D1_(D_z_) and *R*_D3_(3Dz^2^ – 2): 1.4, 3.6, 7.6, 16.6, 33.8, and 50 ms; for *R*_D2_(D_+_): 0.2, 4.7, 9.7, 12.5, 16.5, and 20 ms; for *R*_D4_(D_+_^2^): 3, 6, 12, 22, 34, and 50 ms. All side chain relaxation experiments were done in duplicates. The relaxation data were processed using python scripts, developed based on the scripts used by Hoffmann et al.^46^ Similar to that study, only the original model-free equation (LS2)^39,40^ and LS3 equation (with global tumbling treated as a fitting parameter) were used to fit methyl relaxation data. The global rotational diffusion times of the studied proteins were initially estimated based on backbone *R*_2_/*R*_1_ ratio with the programs R2R1_TM and QUADRIC (www.palmer.hs.columbia.edu).^47^ Global tumbling of all CI2 variants was analyzed with an isotropic diffusion tensor, since D_∥_/D_⊥_< 1.3 for all the variants.

### Chemical shift analysis

We calculated secondary chemical shifts for Cα relative to random coil chemical shifts using the method of Kjaergaard and Poulsen, which corrects for the effect of the neighboring residues (*i*-2, *i*-1, *i*+1, *i*+2, where *i* is the residue of interest) and temperature.^48^

### Entropy calculations using experimental order parameters

Δ*S*^*2*^_*NH*_ and Δ*S*^*2*^_axis_ values obtained in this study were used to estimate the change in conformational entropy upon the mutations by applying the “entropy meter” approach:^49^

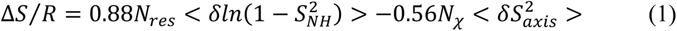

where Δ*S* is the change in conformational entropy, *R* is the gas constant, *N*_*res*_ is the total number of residues, *N*_*χ*_ is the total number of side chain chi angles, 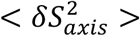 is the measured average change in methyl axis order parameters, and finally 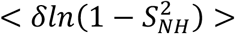 is the corresponding quantity for measured changes in the amide order parameters. The factors of 0.88 and 0.56 account for the reduction in entropy due to correlations of side chain and backbone torsional motions and for the dependence of entropy on order parameters.

### MD simulations

The equations of motion were integrated using the leap-frog integrator implemented in GROMACS v2018.1, using a timestep of 2 fs. We performed the simulations using the Amber a99SB-disp force field with the TIP4P-D-based a99SB-disp water model.^50^ Crystal structures of the CI2 WT (PDB: 7A1H) and its mutated variants L49I (PDB: 7AOK), I57V (PDB: 7A3M) and L49I/I57V (PDB: 7AON) were used as the starting conformations for the simulations.^32^ The proteins were placed in a cubic box of 30.66 nm^3^ (resulting in 0.5 nm distance from the protein to all box edges) and solvated with 6728 to 6836 water molecules. The net charge of CI2 is zero, so no ions were needed to neutralize the system. The system was minimized with 50000 steps of steepest descent and equilibrated for 150 ps in the NPT and NVT ensembles with harmonic position restraints (force constants of 1000 kJ mol^-1^ nm^2^) on the heavy atoms of the protein. Equilibration was performed using velocity rescaling thermostat^51^ with temperature of 298 K, thermostat timescale of 0.1 ps and the Parrinello-Rahman barostat (NPT) with a reference pressure of 1 bar, a barostat time constant 2 ps and an isothermal compressibility of 4.5×10^−5^ bar^-1^. A cut-off range of 1.0 nm was chosen for the Van der Waals and Coulomb interactions and the Particle Mesh Ewald method^52^ was applied to treat long-range electrostatics (4th order cubic interpolation, 0.16 nm grid spacing). The production simulations were run in the NVT ensemble, using velocity rescaling thermostat^51^ with the time constant of 1 ps and the reference temperature of 298 K. We applied harmonic constrains to all bonds involving hydrogens using the LINCS algorithm. All production runs were 1 μs long and the protein coordinates were saved every 1 ps and 3 replicates were used for each simulation, summing to a total of 3 μs per structure. Since the initial structural fluctuations may be related to the equilibration process, all analyses were performed for the last 800 ns of each trajectory, discarding the first 200 ns.

### Calculation of order parameters from MD trajectories

The calculation of the order parameters *S*^*2*^_*NH*_ and *S*^*2*^_*axis*_ was done following the procedures described by Hoffmann et al.^46^ Briefly, we calculated the backbone time correlation function (TCF) of each NH bond vector up to a maximum lag-time of 400 ns. The resulting TCF was then fitted to two different Lipari-Szabo models:

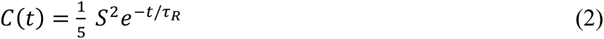

with *S*^2^ used as fitting parameter and

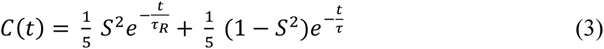

where *τ*^*−1*^ = *τ*_*R*_^*−1*^ + *τ*_*e*_^*−1*^ and with *S*^2^, *τ*_*e*_ as fitting parameters. *C*(*t*) was further multiplied by *ξ* = (1.02/1.041)^6^, to include the effect of vibrational averaging on dipolar couplings.^53^ An isotropic diffusion model was applied using the experimentally derived global rotational diffusion times *τ*_*R*_ (see NMR methods). The obtained order parameters *S*^*2*^_*NH*_ were then averaged over the three independent trajectories. Model selection for each residue was done using the Akaike Information Criterion (AIC):

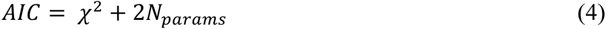

where *χ*^2^ estimates the difference between the fitted and experimental values of the relaxation rates and *N*_*params*_ is the number of the fitting parameters in the model. The model with lowest *AIC* was chosen.

For the side chain order parameters *S*^*2*^_*axis*_, each trajectory was first cut into 10 ns long blocks. After that, internal TCFs (without global tumbling) were calculated for each C_methyl_-H^i^_methyl_ (i = 1,2,3) bond vector of methyl-bearing side chains up to a maximum lag-time of 5 ns and averaged over the three C_methyl_-H^i^_methyl_ (i = 1,2,3) bonds. Next, the internal TCFs were fitted to six exponentials plus an offset,

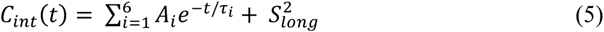

where 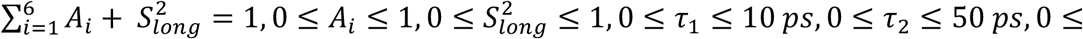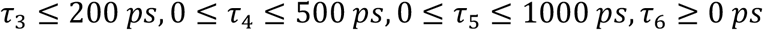 and 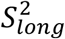 is the long-time limit order parameter. The internal TCF was then multiplied by a single exponential using the experimentally derived global rotational diffusion times *τ*_*R*_:

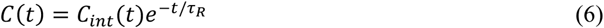

The resulting total TCF was then fitted to

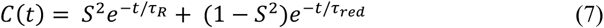

with 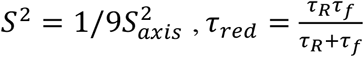 and *S*^2^, *τ*_*f*_ as fitting parameters. Finally, all *S*^2^ where averaged over the 3×80 blocks.

In order to assess the convergence of the calculations we focused on the side chain *S*^*2*^_*axis,sim*_ values, since they, due to the higher flexibility of the side chains, may converge slower than the backbone order parameters.^54–56^ We averaged the 240 original blocks of 10 ns into 10 blocks of 240 ns and performed a leave-N-out cross-validation with *N*=1…5 to evaluate the convergence of the side chain order parameters.^57^ Specifically, for each *N* we split the 10 blocks in a reference set of 10-*N* and a test set of *N* blocks (the left-out blocks), averaged the *S*^*2*^_*axis,sim*_ values within the two sets and computed the RMSD between the resulting average values of the order parameters. For the same *N*, this procedure was repeated for all the possible combinations of the left out blocks and the corresponding RMSDs are averaged to obtain <RMSD>. As shown in **Figure S1** the <RMSD> curves for *N*=5 converge to values around 0.04, which is smaller than the typical difference we computed between the *S*^*2*^_*axis,sim*_ of the different protein variants. For this reason, we can consider our estimations of *S*^*2*^_*axis,sim*_ converged.

### Reweighting simulations using S^2^_*NH*_ and S^2^_*axis*_

The simulated trajectories were analyzed in a block-wise fashion using 100 non-overlapping blocks of 10 ns each. This resulted in 300 blocks for the set of 3x 1 μs-long simulations of each CI2 variant. The backbone *S*^*2*^_*NH*_ and side chain *S*^2^_axis_ order parameters were calculated for each 10 ns block in a similar way as described above but with a maximum lag-time of 5 ns. The resulting TCF was then fitted only to the classical Lipari-Szabo model (equation (5)). Further we performed the weighted average of the order parameters over the blocks, where the weights, **w**, are obtained through the optimization of a functional which determines the agreement between simulated and experimental order parameters. The approach is similar to previously described methods for fitting ensembles using order parameters,^54,58^ but using a reweighting approach similar to our recently described ABSURDer method.^57^ The functional form for the target function that we here used is:

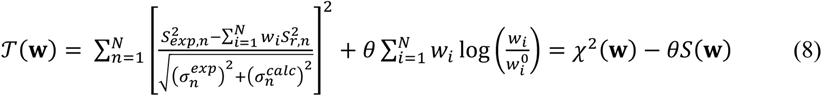

where *n* ranges over the overall number of experimentally defined order parameters and *N* represents the number of blocks which the trajectory has been cut into, 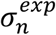 is the experimental error of the *n*-th order parameter, 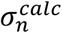 is the standard error of the simulated order parameters, estimated over the different blocks. The optimal set of weights is obtained as the corresponding minimum in weight space, **w**^**\***^ = min_**w**_ *S*^*2*^(*w*). Each weight is associated with a block from the trajectory and it encodes the relevance of each block to the dynamical ensemble. As reference weights we use 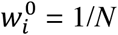 to represent the initial, uniform weights provided by the prior (original simulated ensemble). The parameter *θ* is used to set the balance between our trust in the prior and in the experimental data. See Kümmerer et al.^57^ and Orioli et al.^59^ for the further details.

## Results

Initially, we assessed the overall structural differences in the four variants of CI2 by comparing the chemical shift differences of Cα atoms, in the form of secondary chemical shifts, and of the side chain methyl groups (**Figure 2**). Overall, the changes in chemical shifts reflect only minor and local structural rearrangements upon residue substitutions as expected from the crystal structures of the variants.^32^

**Figure 2.**
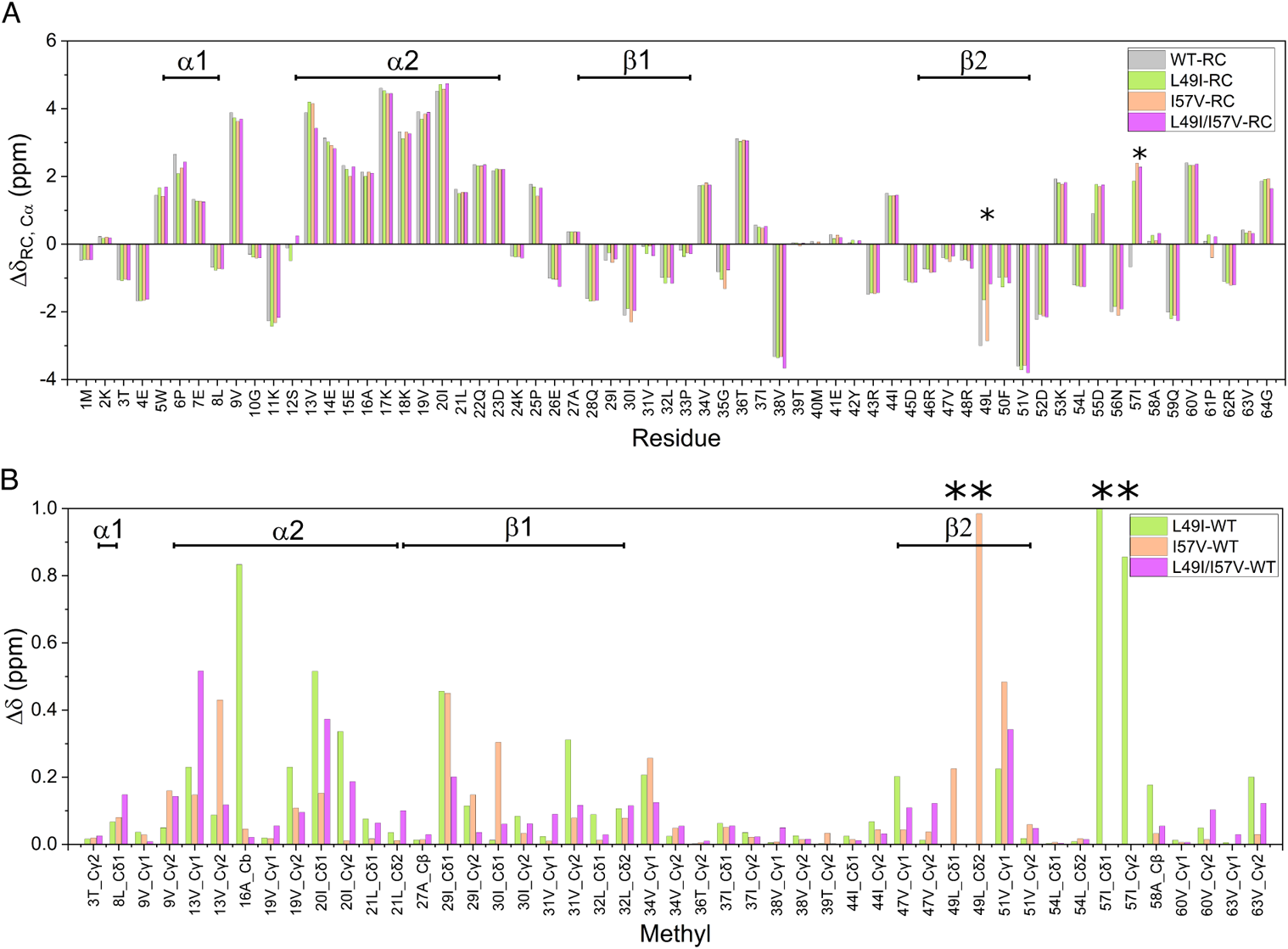
Chemical shift analysis. (A) Secondary Cα chemical shifts. (B) Absolute difference in methyl ^13^C CSs of the wild type (WT) and mutated variants of CI2. Mutation sites are highlighted with the asterisks. Secondary structure elements are depicted at the top of each panel.

### Backbone dynamics

We assessed the conformational dynamics on the ps-ns timescale of the four CI2 variants using NMR relaxation experiments and interpreted it with the model-free (MF) formalism. We analyzed both the dynamics of the backbone amide groups and of the side-chain methyl groups for Ala, Ile, Leu, Thr, and Val. The side-chain methyls of the two Met residues were excluded from the analysis due to low data quality.

For the backbone ^15^N amides, we acquired *R*_1_, *R*_2_ and {^1^H}-^15^N NOE relaxation parameters at two magnetic fields (600 and 750 MHz) for each residue. The six experimental data points allowed us to sample the spectral densities, *J*(ω), at five different frequencies for each residue. We analyzed the data with the model-free formalism and to avoid overfitting we excluded MF models containing more than three parameters. Since the initial analysis in *relax* showed that the ratio between parallel and perpendicular components (D_∥_/D_⊥_) of the rotational diffusion tensor was less than 1.3 for all four CI2 variants, we used an isotropic model for the global molecular tumbling. For each CI2 variant, we were able to analyze the data from all non-proline backbone amides, except for K2. The consistency of the relaxation data for the remaining residues were assessed by comparing the values of the spectral densities at zero frequency, *J*(0), calculated from data collected at either of two different fields. In the absence of μs–ms motions, the *J*(0) values calculated from the two datasets are expected to be the same.^60^ For all four CI2 variants *J*(0) values from the two fields are highly correlated (**Figure S2**). Most of the deviating amide groups are also characterized by elevated *R*_1_*R*_2_ values. Unlike *R*_2_/*R*_1_ ratio, the *R*_1_*R*_2_ product includes the effects of chemical exchange but not the effects of motional anisotropy.^61^ Comparison of the *R*_1_*R*_2_ values between the CI2 variants suggests that L49I may experience a decrease in μs-ms conformational dynamics whereas I57V and L49I/I57V, oppositely, may experience an increase compared to the wild type (**Figure S3**). We note that a high correlation between the *R*_2_/*R*_1_ ratio and the *R*_1_*R*_2_ product suggests little motional anisotropy and thus justifies an isotropic model for the global molecular tumbling of CI2 variants.

Overall, the order parameters, *S*^2^_*NH*_, for the backbone amides are quite similar for all variants with a value around 0.8 in the structured regions of the protein. which drops to ~0.7 in the more flexible inhibitory loop around residues I37-I44 (**Figures S4A**). The lowest value, indicating greatest flexibility, is found at residue M40 that plays an important functional role in chymotrypsin inhibition. We note that several residues around this region, according to their elevated *R*_1_*R*_2_, values experience slower dynamics.

We compared the *S*^2^_*NH*_ values between the variants of CI2 to assess the effects of the mutations on the dynamics. To minimize the effects of outliers, we calculated the Δ*S*^2^_*NH*_ values as the difference between the *S*^2^_*NH*_ values for each variant and the average values of the other three variants, as recently suggested.^62^ For example, in the case of WT: Δ*S*^2^_WT_ = *S*^2^_WT_ – <*S*^2^>_L49I, I57V, L49I/I57V_. The analysis demonstrates that the differences in ps-ns conformational dynamics overall is small, however, we do see some sequence dependent variation (**Figure 3**). Δ*S*^2^_WT_ is positive for the first eleven residues, residues 26E, 27A, and in the region close to the mutation site 57 (53K-56N, and 59Q-60V) suggesting smaller than average amplitude of the backbone dynamics of WT for these residues. For most of the other residues Δ*S*^2^_WT_ are around 0.01-0.02 units lower than the average values of the other three variants. The destabilizing L49I mutation results in a decrease in the dynamics of almost all amide groups in CI2: most of ΔS^2^_L49I_ values are in between 0.01-0.2 units. The stabilizing I57V mutation results in a distribution of Δ*S*^2^_I57V_ values which almost mirrors that for WT, namely positive Δ*S*^2^_I57V_ for the first eleven residues and for residues 26E, 27A, 53K-56N, and 59Q-60V. The I57V/L49I variant is characterized by the increased dynamics for almost all backbone amide groups. The summed (excluding the mutation sites) differences in the backbone order parameters, 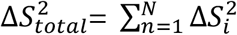 (where *i* is an amide group and *N* is the total number of amide groups), are listed in **Table 1**.

**Table 1.**
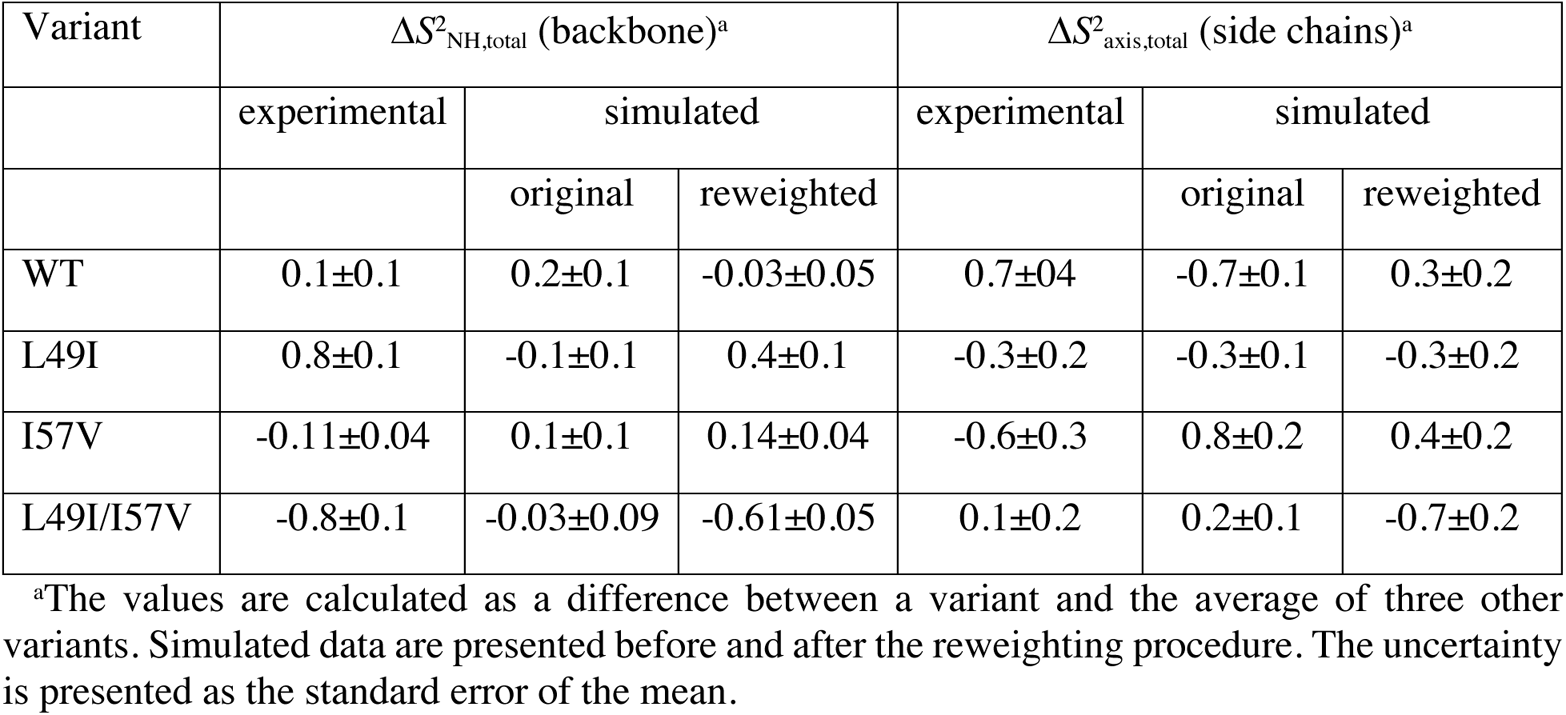
The total average difference in order parameters for each CI2 variant.

| Variant | $\Delta S^2_{NH, total}$ (backbone)<sup>a</sup> | | | $\Delta S^2_{axis, total}$ (side chains)<sup>a</sup> | | |
| --- | --- | --- | --- | --- | --- | --- |
|  | experimental | simulated |  | experimental | simulated |  |
|  |  | original | reweighted |  | original | reweighted |
| WT | 0.1±0.1 | 0.2±0.1 | -0.03±0.05 | 0.7±0.4 | -0.7±0.1 | 0.3±0.2 |
| L49I | 0.8±0.1 | -0.1±0.1 | 0.4±0.1 | -0.3±0.2 | -0.3±0.1 | -0.3±0.2 |
| I57V | -0.11±0.04 | 0.1±0.1 | 0.14±0.04 | -0.6±0.3 | 0.8±0.2 | 0.4±0.2 |
| L49I/I57V | -0.8±0.1 | -0.03±0.09 | -0.61±0.05 | 0.1±0.2 | 0.2±0.1 | -0.7±0.2 |

<sup>a</sup>The values are calculated as a difference between a variant and the average of three other variants. Simulated data are presented before and after the reweighting procedure. The uncertainty is presented as the standard error of the mean.

**Figure 3.**
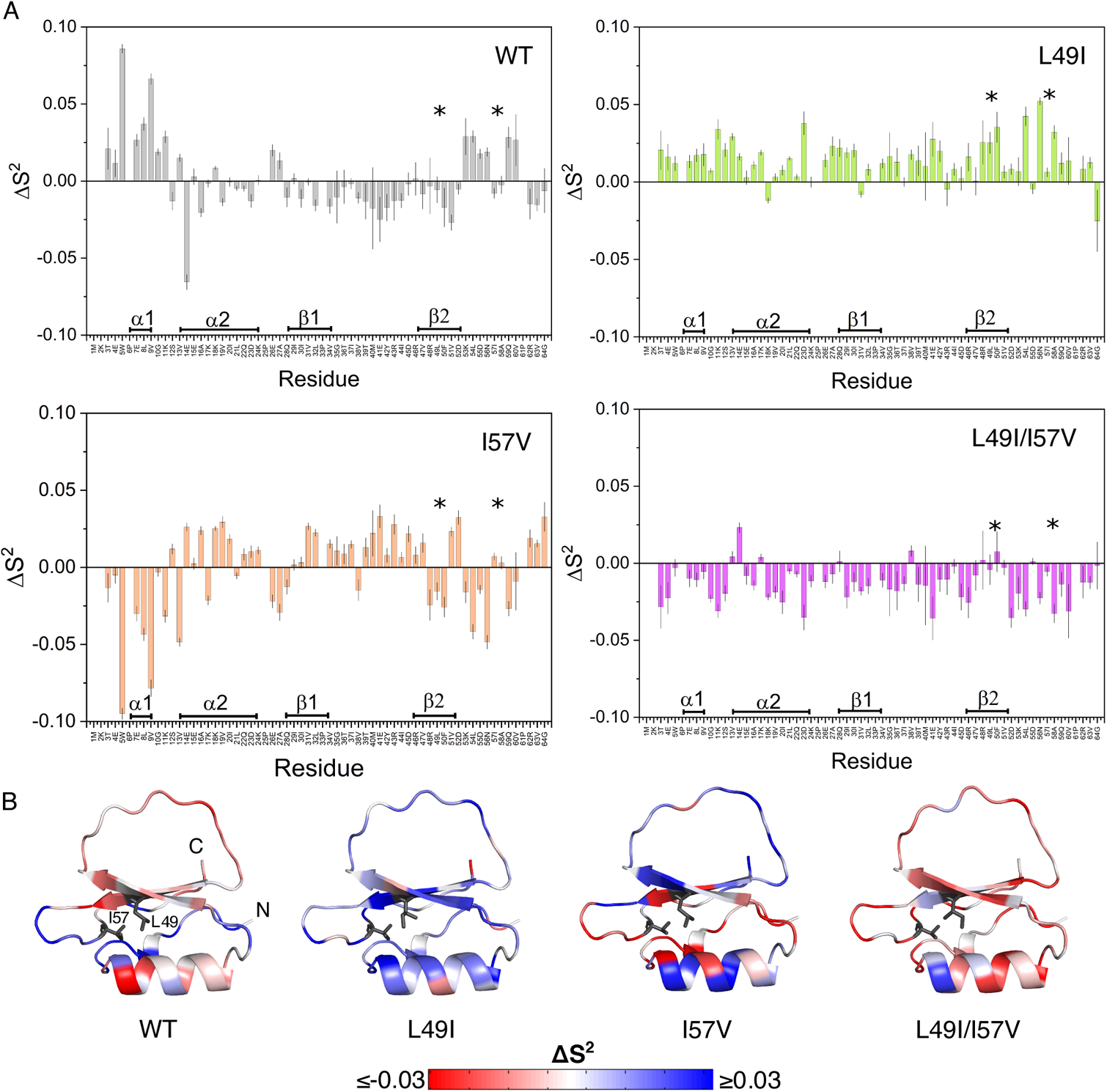
The difference in NMR-derived order parameters (for backbone N-H groups) between one of the CI2 variants and the average value of three other CI2 variants: ΔS^2^_*NH*_ = S^2^_*NH*_(variant) – S^2^_*NH*_ (average-of-three-other-variants). Secondary structure elements are shown at the bottom of each panel. Mutation sites are highlighted by asterisks. (A) ΔS^2^_*NH*_ plots (B) The difference in order parameters is shown as a color scale on the structure of CI2.

### Side chain dynamics

We assigned the chemical shifts for all side chain methyl groups of WT CI2. However, some were excluded from the relaxation analysis. For L8 ^2^H-C^δ2^, V31 ^2^H-C^γ1^, and V63 ^2^H-C^γ1^ it was not possible to estimate the volume or the maximum height due to significant peak overlap. Moreover, the chemical shifts for L8-C^δ1^, L32-C^δ1^, and L49-C^δ1^ (in case of WT and I57V) were greater than 24 ppm. This may result in strong ^13^C–^13^C coupling between C^γ^ and C^δ^ in leucine residues and the ^2^H relaxation data can be compromised.^46^ In total, we included 42 peaks in the analysis of side chain ps-ns dynamics.

Fast dynamics of side chain CH_2_D isotopomers can be tracked based on two to five relaxation rates. In this study we acquired four side-chain relaxation rates at two magnetic fields (600 and 750 MHz):^44,45^ *R*_D1_(D_z_) – longitudinal magnetization; *R*_D2_(D_+_) – in-phase transverse magnetization; *R*_D3_(3Dz^2^ – 2) – quadrupolar order magnetization; and *R*_D4_(D_+_^2^) – double quantum magnetization. Collection of these four relaxation rates results in an overdetermination since they depend on the spectral densities at only three different frequencies (zero frequency *J*(0), the deuterium frequency at the given field strength *J*(ω_D_), and double frequency of deuterium *J*(2ω_D_)).^44^ Therefore, according to the relationship *R*_D4_ = 1/2*R*_D1_ + 1/6*R*_D3_ we found that the consistency of the datasets at both magnetic fields is good (R^2^ ≥ 0.97, **Figure S5**). All acquired side chain deuterium relaxation rates were fitted to LS2 and LS3 model-free models.

As in case of backbone amide order parameters, side chain methyl *S*^2^_axis_ values are quite similar for all the variants (**Figure S4B**). Similar to *S*^2^_NH_, we calculated Δ*S*^2^_axis_ values as the difference between *S*^2^_axis_ values of each of the variants and the average values of other three variants (**Figure 4**). A direct comparison suggests that there may be subtle changes in the ps-ns dynamics of side-chain methyl groups. However, the errors are close to the value of Δ*S*^2^_axis_. In general, according to the total summed Δ*S*^2^_axis_, side chain methyl groups of the three mutants are more flexible than the wild type (**Table 1**).

**Figure 4.**
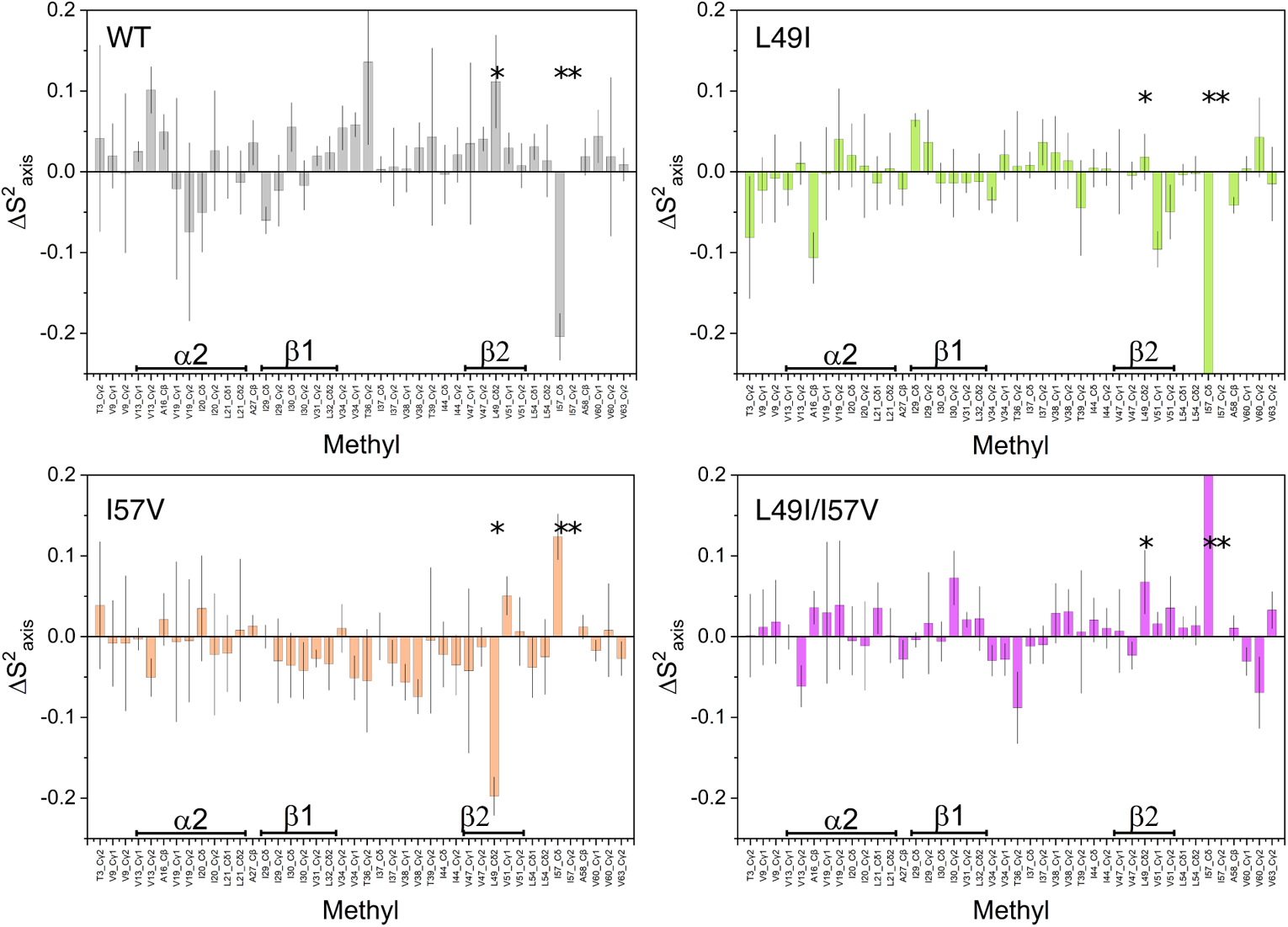
The difference in NMR-derived squared generalized order parameters (for sidechain methyl groups) between one of the CI2 variants and the average value of three other CI2 variants: ΔS^2^_axis_ = S^2^_axis_(variant) − S^2^_axis_(average-of-three-other-variants). The borders of the structural elements are depicted at the bottom of each panel, mutation sites are highlighted by asterisks.

### Entropy calculations

To obtain a quantitative measure of the effects of the mutations we used the Δ*S*^2^_NH_ and Δ*S*^2^_axis_ values to estimate the change in the conformational entropy by applying the “entropy meter” approach.^63,49^ The entropy contribution to the free energy calculated in this way correlates well with experimental ΔΔ*G*_f_ values (Pearson’s *r* = −0.96; **Figure 5**). The individual contributions from the backbone and the side-chains are in the same order of magnitude although most of the error in the total value originate from the side-chain data (**Table S1**). We note that the variation in the calculated TΔ*S* is larger than the variation in ΔΔ*G*_f_, which could either result from entropy-enthalpy compensation or because the “entropy meter” overestimates the entropy in our case. Still, the conclusion from the strong correlation is that changes in conformational entropy are important for the changes in conformational stability we observe for CI2.

**Figure 5.**
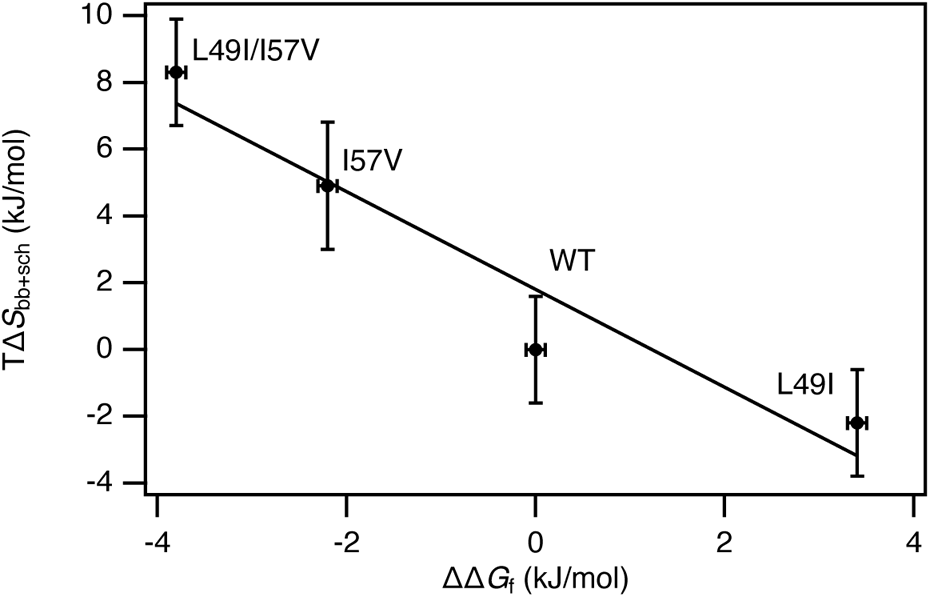
Correlation between changes in thermodynamic stability of CI2 variants values from unfolding experiments (ΔΔ*G*_f_) and changes in the entropy contribution from ps-ns dynamics to the free energy (TΔ*S*_bb+sch_).

### Molecular dynamics simulations

We performed MD simulations of the CI2 variants using a99SB-disp with the modified TIP4PD water model,^50^ a force field that substantially improves the accuracy for simulations of the disordered elements without sacrificing accuracy for structured regions. This force field was chosen due to the possible higher flexibility of several residues in the inhibitory CI2 loop. For all CI2 variants we conducted 3 simulations of 1 μs each, obtaining a total of 3 μs of sampling per variant.

As two rotamers of I49 are observed in the crystal structure of the L49I/I57V variant (PDB: 7AON) we ran two independent simulations starting from each of the rotamers. In the first rotamer structure, l49I/I57V(A), the I49 χ^1^ angle (N–Cα–Cβ–Cγ) is in gauche-conformation and in the second structure, l49I/I57V(B), the I49 χ^1^ angle is in gauche+ conformation. For L49I/I57V, the I49 χ^1^ angle populates the gauche-rotamer for 95% of the time and the gauche+ for the remaining 5%. This behavior is independent of the initial rotameric state. I49 χ^1^ flips from gauche+ to gauche-within the first 15 ns of the simulations, which indicated that the force field prefers the gauche-conformation of I49 χ^1^. We therefore only consider l49I/I57V(A) in the following. Initial analysis of the simulated ensembles revealed no changes in the overall structure of either of the variants: the average all-atom RMSD is in the range of 0.26-0.29 nm and per-residue RMSF does not exceed 0.6 nm for the most flexible residue (M40) in all five systems.

### Simulation-derived order parameters

For a more detailed comparison of NMR and MD data we analyzed the acquired trajectories in terms of backbone amide and side chain methyl order parameters. As for the experimental data we used the model-free formalism. For consistency with experimental data the extended model-free models were omitted. The impact of the amount of sampling on the values of the order parameters was checked by employing a leave-N-out cross-validation. The analysis suggests that our estimations of the order parameters are converged (**Figure S1**).

*S*^*2*^_*NH*,sim_ values are quite similar for all CI2 variants and reflect the absence of any major effect of the mutations. However, the correlations between *S*^*2*^_*NH*,sim_ and *S*^*2*^_*NH*_ for the four variants are not strong (Pearson’s correlation coefficient, *r* = 0.55-0.62; **Figures S6A**). This may be related to several issues. First, the simulations may not be converged, though our analysis described above suggests this to be a minor effect. Second, the ^15^N transverse relaxation rates can be significantly affected by μs-ms timescale dynamics itself (*R*_ex_ contribution, **Figure S3**) and it is not always easy to accurately correct for these effects.^64,65^ Third, despite the application of a state-of-the-art force field, imperfections can still reduce the match between simulations and experiments. These limitations of the computational approach may significantly affect not only the absolute values of the order parameters but also the difference in the dynamics of CI2 variants. As in case of the backbone, limited agreement between NMR and MD sidechain order parameters (*S*^2^_axis,sim_) may be related to the force field imperfections and was observed in other studies (**Figure S6B**).^46,66^

### Reweighting MD trajectories

It was shown previously that the direct integration of experimental data with molecular simulations can significantly improve a detailed description of the structural and dynamic properties of biomolecules. Such integration can be performed *a priori* using MD simulations biased with experimental data or by reweighting unbiased simulations using experimental data *a posteriori*.^67^ The second approach allows us to reweight the simulated trajectories also using our NMR relaxation data.^59^ The relaxation rates are used in the fitting procedure to obtain *S*^2^_*NH*_ and *S*^2^_*axis*_ and can be considered as a source of experimental data.^46,66^ Due to slow μs-ms time scale motions for some residues in CI2, the *R*_2_ rate values can be affected by exchange (**Figure S3**). This contribution is not captured in MD simulations, which can result in overfitting during the reweighting procedure if we directly target the relaxation rates. Consequently, instead we reweighted our simulated trajectories using *S*^2^_*NH*_ and *S*^2^_*axis*_ values. We used an approach inspired by our recently developed ABSURDer method^57^ but targeting the order parameters as also done previously.^54,58^ For each CI2 variant *S*^2^_*NH*,sim_ and *S*^2^_*axis*,sim_ values were calculated separately for each of the 300 blocks of 10 ns and further reweighted against the experimental *S*^2^_*NH*_ and *S*^2^_*axis*_ values.

The reweighting procedure resulted in a noticeable improvement of the agreement between the experimental and computational data (**Figure 6**). Pearson’s correlation coefficient (*r*) and RMSD for the backbone *S*^*2*^_sim_ and *S*^*2*^_*NH*_ improves for all CI2 variants (from *r* = 0.45-0.62, RMSD = 0.05- to *r* = 0.59-0.71, RMSD = 0.04-0.05; **Figures 6A**). Similar to the experimental data, we compared Δ*S*^2^_*NH*,sim_ values calculated as a difference between *S*^*2*^_*NH*,sim_ values of each of the variants and the average values of other three variants.^62^ The Δ*S*^2^_*NH*,sim,total_ values get closer to the experimental Δ*S*^2^_*NH*,total_ values and better reflect experimentally observed changes in the order parameters (with an exception of I57V; **Table 1**). In case of the side chain methyl order parameters (*S*^*2*^_axis,sim_) after the reweighting *r* and RMSD also improve (from *r* = 0.55-0.69, RMSD = 0.27-0.30 to *r* = 0.62-0.87, RMSD = 0.12-0.27; **Figure 6B)**.

**Figure 6.**
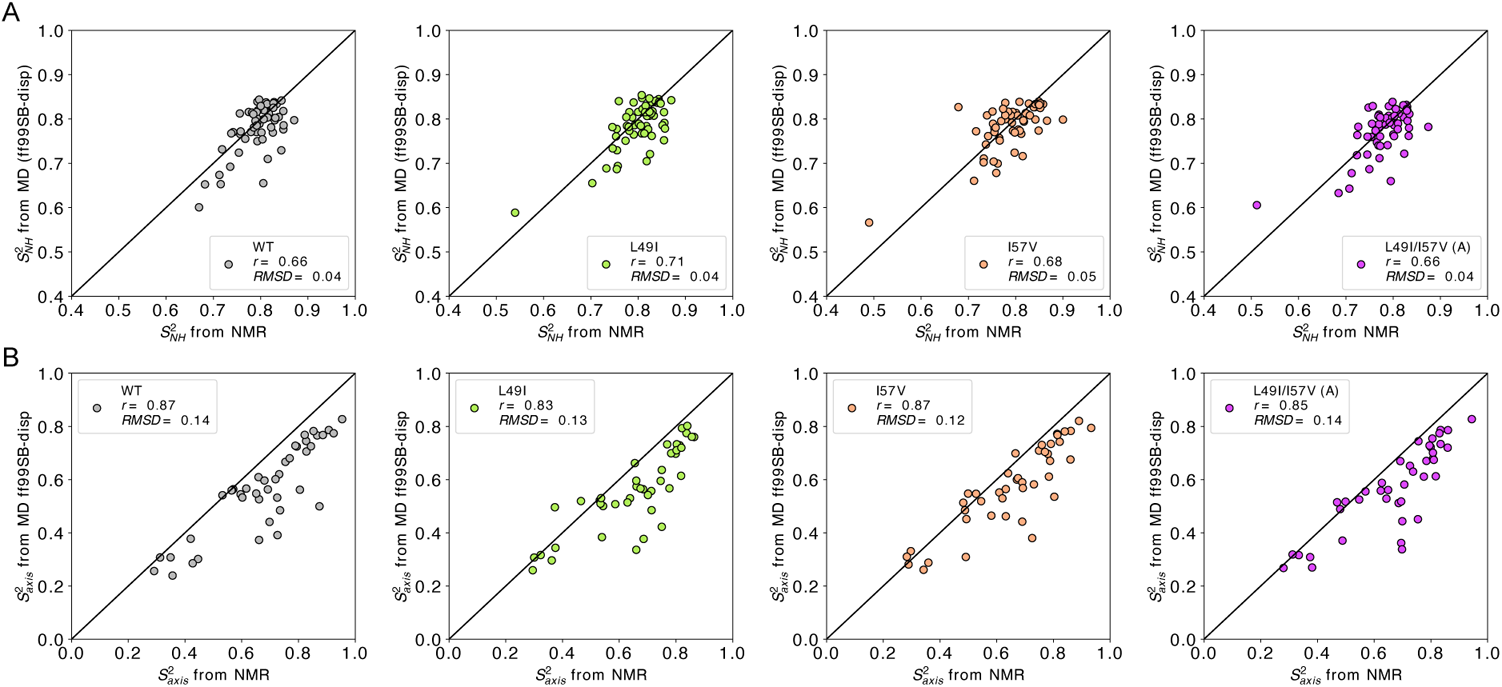
Experimental backbone *S*^2^_*NH*_ (A) and side chain *S*^2^_axis_ (B) order parameters of CI2 compared to their ***reweighted*** computational counterparts extracted from MD simulations. Pearson correlations (*r*) and RMSDs were calculated for each variant separately.

The trajectory reweighting results in a significant improvement of the correlation between experimentally measured and MD-derived backbone and/or side chain order parameters for all CI2 variants, except I57V. However, even after the reweighting the MD simulations do not capture all small variations in the dynamics of the four CI2 variants. Consequently, we omitted calculations of conformational entropy from the simulations. Similar to the analysis of the experimental data (**Figures 3 and 4**), the reweighted computational ensembles did not reveal any clear changes in the structural and dynamic properties of CI2. This suggests that the changes in stability and conformational entropy are the combined effect of several subtle contributions. In accordance with the previous study of CI2 dynamics,^26^ the mutations thus affect the conformational dynamics of all 64 amino acid residues as a whole, probably by introducing many small changes in the contact network of the protein.

## Discussion

We have demonstrated that ps-ns conformational dynamics of CI2 can be affected by mutations in the hydrophobic core and find that the level of the conformational entropy correlates with ΔΔ*G*_f_. Based on the NMR measurements, we conclude that the entropy and in particular the backbone entropy provides a major contribution to the stability of the CI2 variants analyzed in this work. From the entropy meter estimates the increase in entropy is even larger than the gain in conformational stability suggesting that the mutations could be associated with an unfavorable loss of intermolecular interactions. We note, however, complications arising from entropic contributions arising from dynamics on time scales not probed by the relaxation measurements,^68^ and which may not be accurately captured by the entropy meter approach. Our analysis of the crystal structures of the four CI2 variants showed that the structures are very similar and only subtle structural changes are observed throughout the protein.^32^ This picture is mirrored by our current NMR analysis. The very small changes in dynamics between the variants are thus scattered throughout the protein. The largest difference is near the N-terminal for residues 5-9 that form a helical turn. This region appears most rigid in the wild type and most dynamic in the I57V variant. We speculate that the packing between the residue at position 57 and V9 is slightly weakened by the presence of a Val position 59 and that this effect is alleviated by having an Ile at position 49. However, we do not see any structural rearrangements around V9 or any other residues in the helical turn.

Our MD simulations of the CI2 variants support our findings from the experimental data, i.e. that differences in the dynamics are subtle and distributed throughout the protein, although the simulations cannot fully reproduce our experimental data. The difference between NMR and MD derived dynamic properties may be related to the presence of fluctuations in the native state of CI2 and its variants that can affect fast protein dynamics but cannot be intensively sampled by relatively short simulations. This assumption may be supported by a previous computational study of CI2.^69^ In that work MD simulations, restrained with protection factors for local H-exchange, revealed a native state ensemble of CI2 including rare, large fluctuations from the native state, which may take place on a timescale of milliseconds or more and cannot be related to the folding pathway of CI2.^69^ This is consistent with our experimental *R*_1_*R*_2_ values suggesting that mutations of CI2 may also affect the dynamics on slower (μs-ms) time scales.

Our best explanation for increased stability is thus that the mutations releases the previously proposed unfavorable strain in the hydrophobic mini-core of A16, L49 and I57,^70^ and that this occurs without any actual rearrangement of the structure and thus of the enthalpic interactions that stabilize the folded state. This is much in line with the conclusions based on a structural analysis of the I57V mutant by Fersht and co-workers.^71^

In summary, we have demonstrated that in case of CI2 and its mutated variants, changes in fast ps-ns dynamics may significantly contribute to changes in protein stability. Our analysis suggests that also slower μs-ms timescale dynamics of CI2 may be affected by these mutations.

## AUTHOR INFORMATION

### Notes

The authors declare no competing financial interest.

## ACKNOWLEDGMENT

The authors would like to thank Falk Hoffmann for the help with the calculation of order parameters from MD simulations and Yong Wang for the assistance with MD simulations setup.

