## Supporting Figures and Tables for "Double mutant of chymotrypsin inhibitor 2 stabilized through increased conformational entropy"

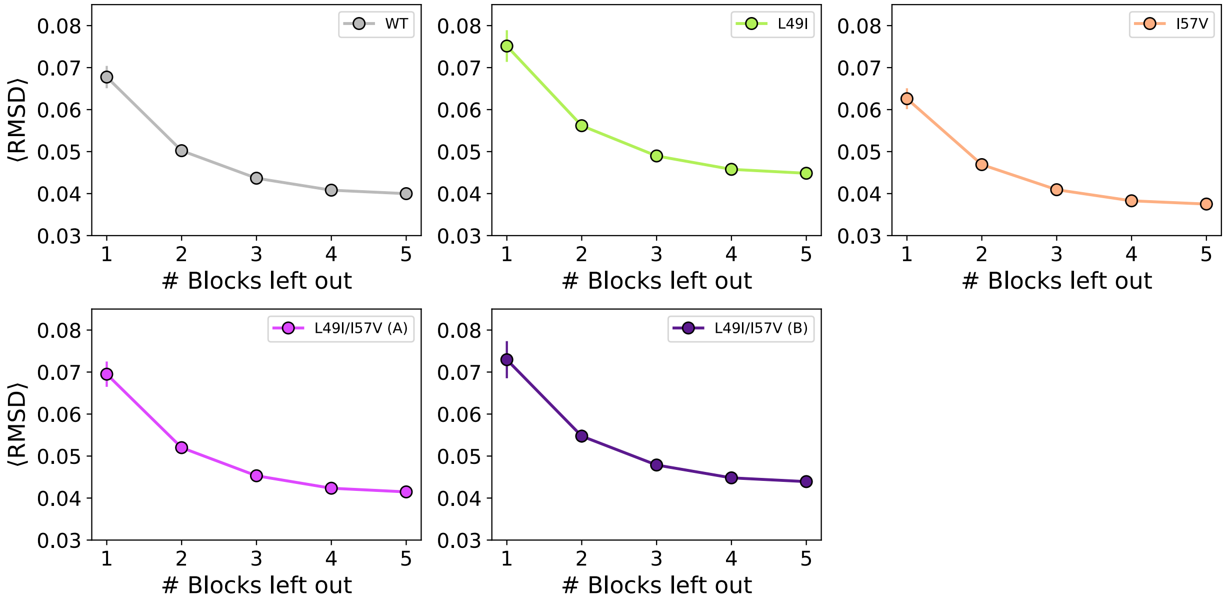


**Figure S1.** Convergence of MD derived side chain order parameters ($S^2_{axis,\ sim}$) using Amber ff99SB-disp force field.

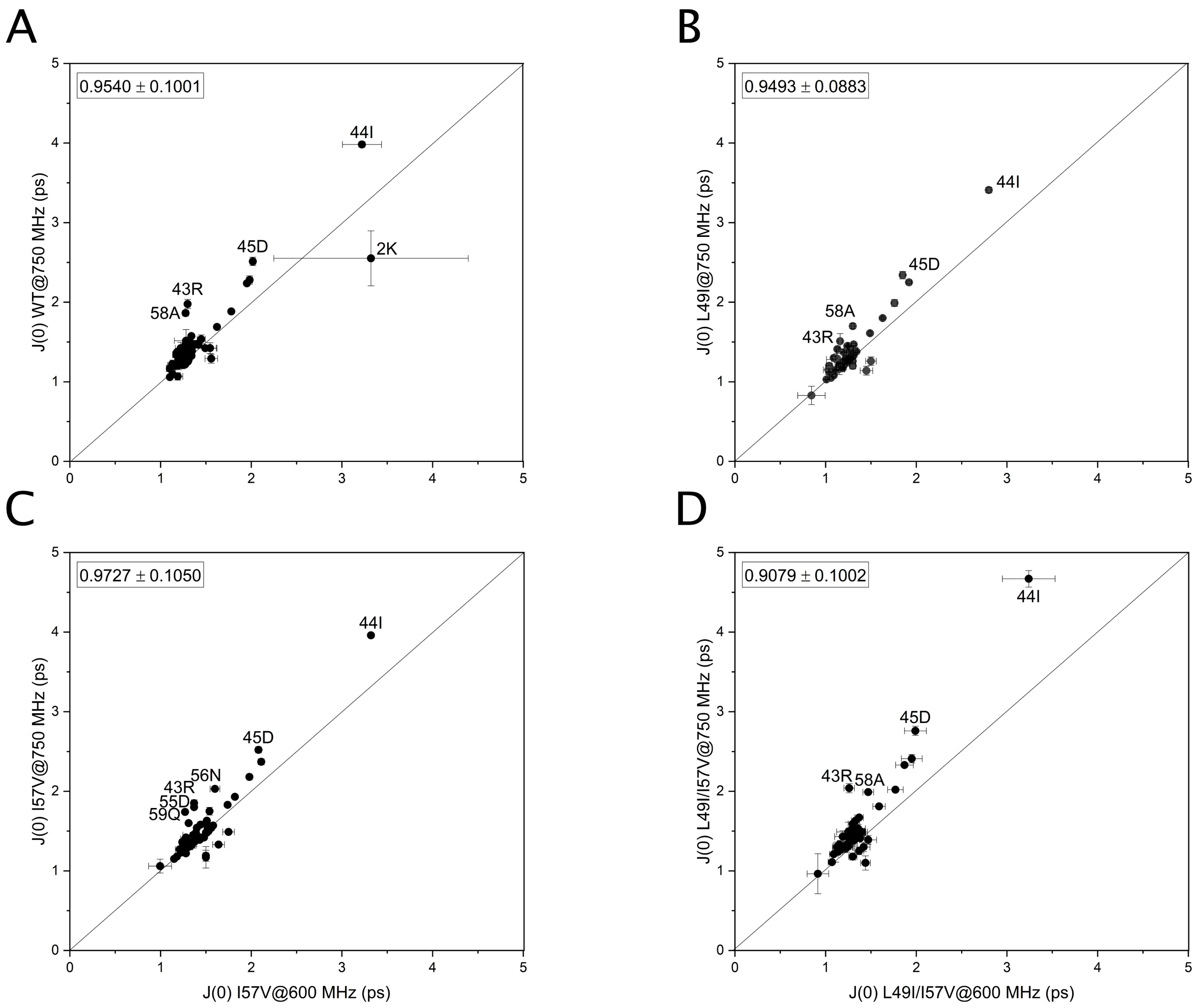


**Figure S2.** Consistency of calculated spectral densities. (A) WT; (B) L49I; (C) I57V; (D) L49I/I57V double mutant. Cells at the top of the plots represent the average value and standard deviation of ratio between 600 and 750 MHz data. Amide groups with the highest difference between 600 and 750 MHz data are labeled. The consistency of two magnetic field datasets is high. Most of the deviating amide groups also have elevated $R_1R_2$ values and are expected to experience slow (μs-ms) dynamics.

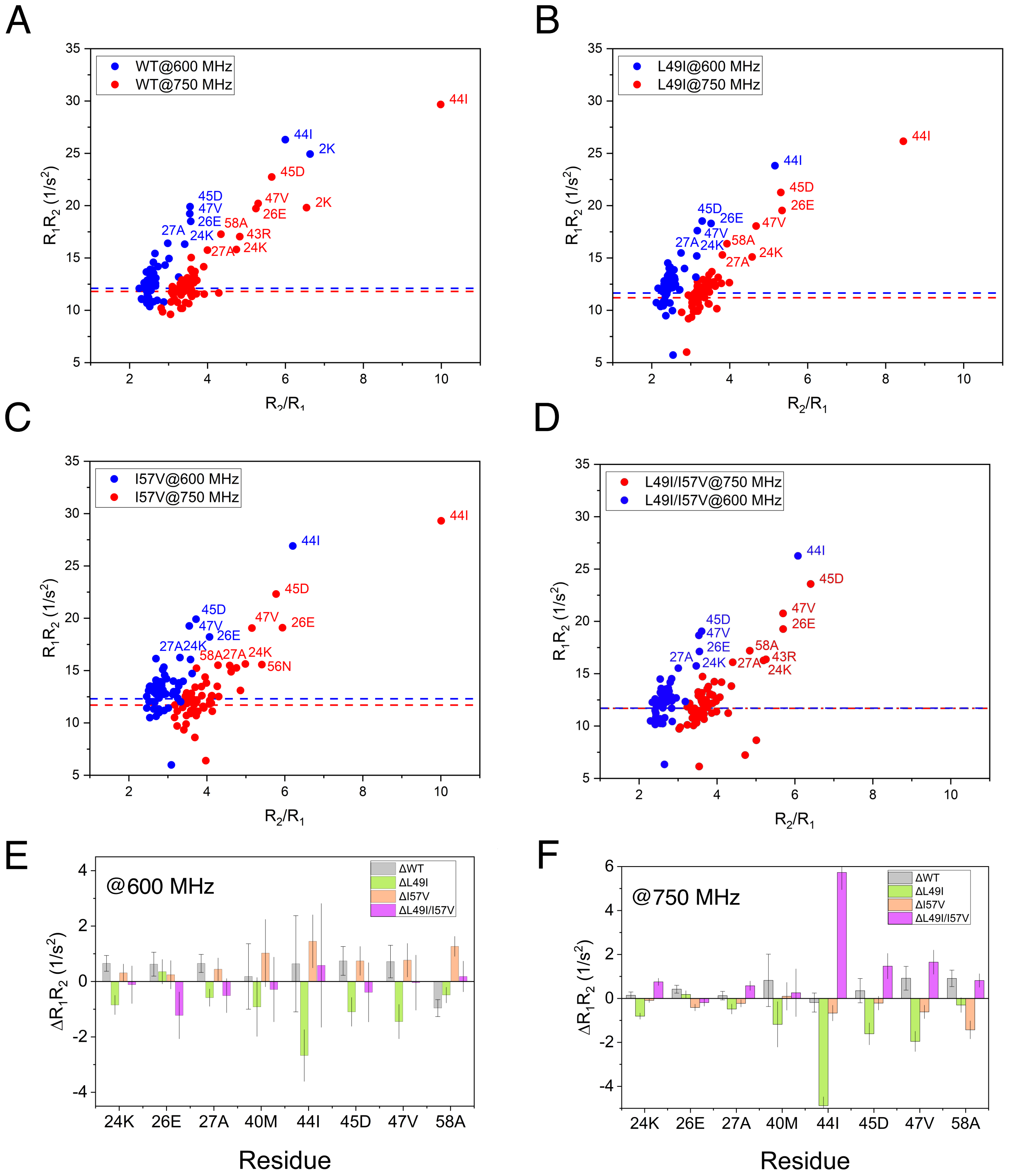


**Figure S3.** Exchange outliers. (A-D) $R_2/R_1$ vs $R_1R_2$ plots for the backbone relaxation rate of CI2 variants. (A) WT; (B) L49I; (C) I57V; (D) L49I/I57V double mutant. The values are present for the data acquired at two fields (600 and 750 MHz). Backbone amides with elevated $R_1R_2$ values are labeled; 10% trimmed mean values after exclusion of residues with low NOE values for both fields are highlighted with the dashed lines. (E-F) The difference in $R_1R_2$ values between one of CI2 variants and the average value of three other CI2 variants for the labeled amides from A-D ($\Delta R_1R_2$). The data acquired at 600 MHz (E) and 750 MHz (F).

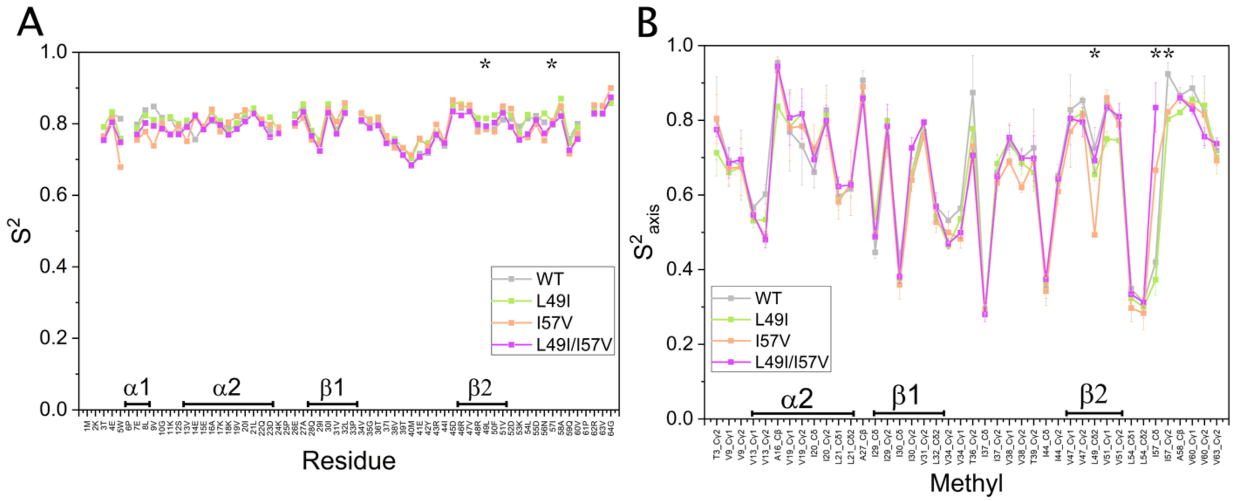


**Figure S4.** NMR-derived squared generalized order parameters for: (A) Backbone amide groups ($S^2$); (B) Side-chain methyl groups ($S^2_{axis}$). The borders of the structural elements are depicted at the bottom of each section, mutation sites are highlighted with the asterisks.

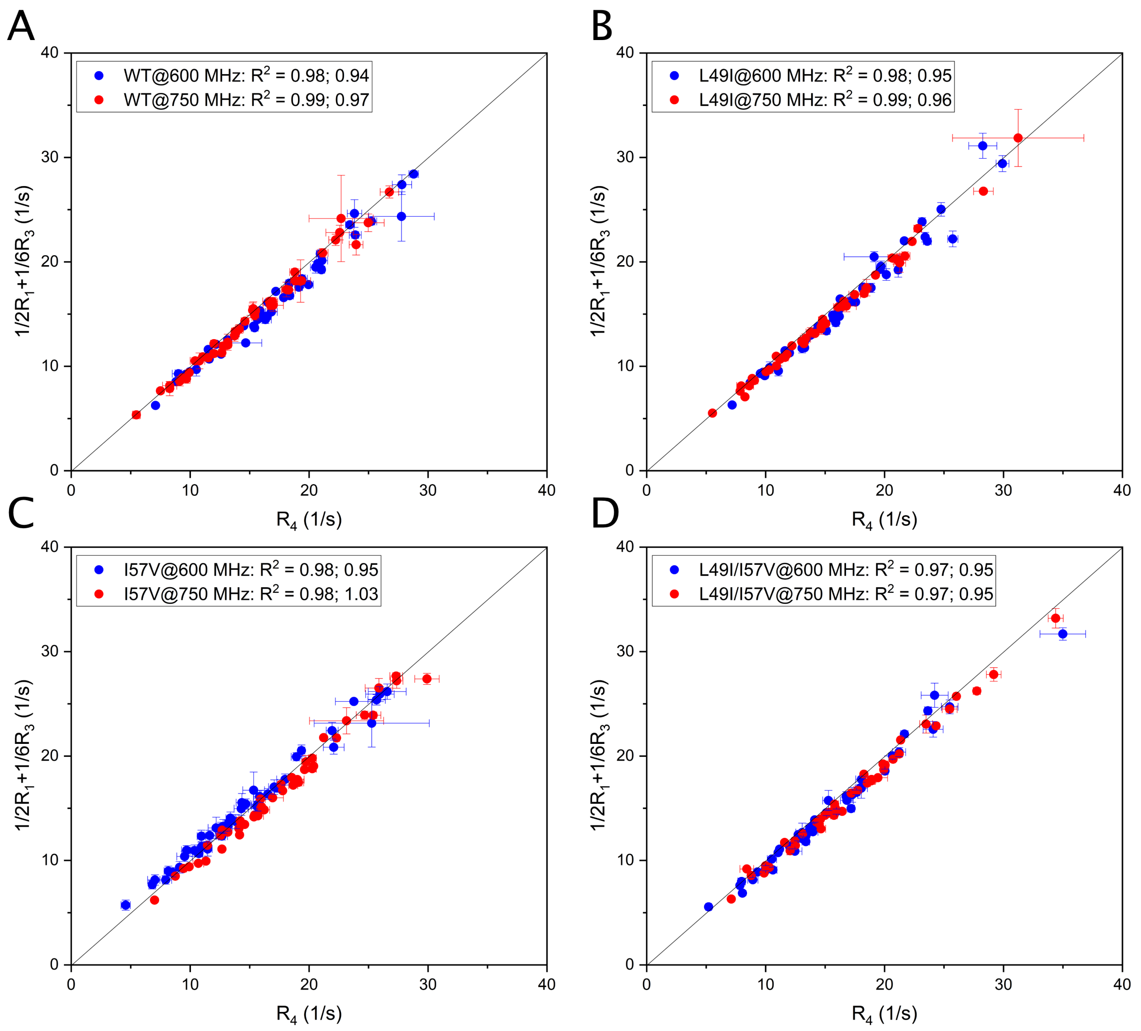


**Figure S5.** Consistency of datasets for CI2 variants collected at 600 and 750 MHz. (A) WT; (B) L49I; (C) I57V; (D) L49I/I57V double mutant. The correlation is based on the ratio: $R_{D4} = 1/2R_{D1} + 1/6R_{D3}$. The correlation plots with a diagonal at $x = y$. The consistency of the datasets at both magnetic fields is high. The numbers in the legend: $R^2$ correlation coefficient and the average value of the ratio between $R_{D4}$ and $[1/2R_{D1} + 1/6R_{D3}]$ values.

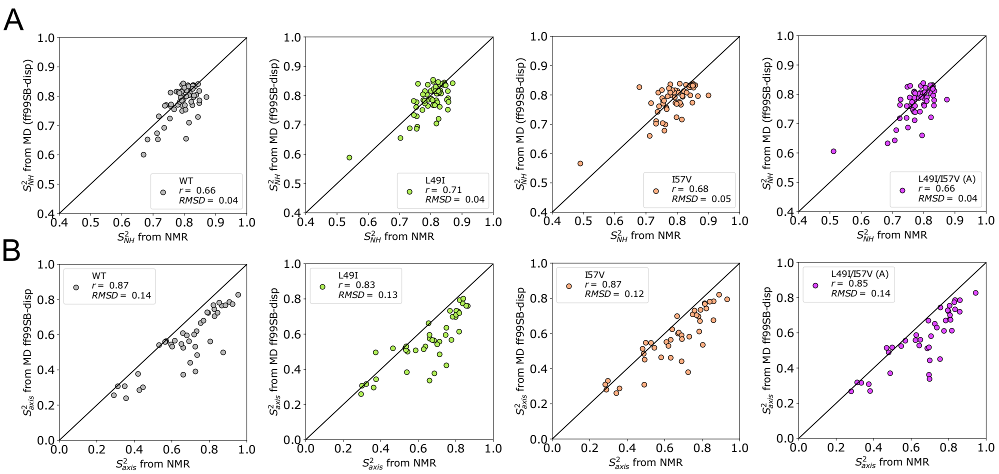


**Figure S6.** Backbone $S^2_{\mathrm{NH}}$ (A) and side chain $S^2_{\mathrm{axis}}$ (B) order parameters of CI2 from ***reweighted*** MD and experiments extracted from Lipari-Szabo (LS) model free fitting. Scatter plots of $S^2_{\mathrm{NH}}$ and $S^2_{\mathrm{axis}}$ from NMR and MD using Amber ff99SB-disp force field. Pearson correlations (*r*) and RMSDs were calculated for each variant separately.

**Table S1.** Entropy changes from entropy-meter calculation.

| | $T\Delta S_{bb}$ | $T\Delta S_{sch}$ | $T\Delta S_{bb+sch}$ |
|---|---|---|---|
| L49I-WT | -5.5±0.2 | 3.3±1.6 | -2.2±1.6 |
| I57V-WT | 0.6±0.2 | 4.3±1.9 | 4.9±1.9 |
| L49I/I57V-WT | 6.3±0.2 | 2.0±1.6 | 8.3±1.6 |

The total average difference between a mutant and WT backbone ($T\Delta S_{bb}$), sidechain ($T\Delta S_{sch}$) and total ($T\Delta S_{bb+sch}$) conformational entropy contribution to the free energy (kJ/mol). The uncertainty is presented as the standard error of the mean.
